# Transposable elements continuously remodel the regulatory landscape, transcriptome, and function of decidual stromal cells

**DOI:** 10.1101/2021.10.25.465769

**Authors:** Katelyn Mika, Vincent J. Lynch

## Abstract

Gene expression evolution underlies the origin, divergence, and conservation of biological characters including cell-types, tissues, and organ systems. Previously we showed that large-scale gene expression changes in decidual stromal cells contributed to the origins of pregnancy in eutherians and the divergence of pregnancy traits in primates, and that transposable elements likely contributed to these gene expression changes. Here we show that two large waves of TEs remodeled the transcriptome and regulatory landscape of decidual stromal cells, including a major wave in primates. Genes nearby TE-derived regulatory elements are among the most progesterone responsive in the genome and play essential roles in orchestrating progesterone responsiveness and the core function of decidual cells by donating progesterone receptor binding sites to the genome. We tested the regulatory abilities of 89 TE consensus sequences and found nearly all acted as repressors in mammalian cells, but treatment with a histone deacetylase inhibitor unmasked latent enhancer functions. These data indicate TEs have played an important role in the development, evolution, and function of primate decidual stromal cells and suggest a two-step model in which latent enhancer functions of TEs are unmasked after they lose primary repressors functions.

**Significance statement:** Gene expression patterns evolved very rapidly during the evolutionary origins of pregnancy in early mammals and in primates. These episodes of gene expression evolution and linked with important processes that establish and maintain pregnancy, and appear to be driven by domestication of transposable elements.

## Introduction

Changes in gene regulatory evolution underlie the origin, divergence, and homology of cell-types, tissues, and organs. Thus, understanding the mechanisms of gene regulatory evolution is essential for understanding how anatomical systems evolve. One hypothesis of gene regulatory evolution proposes that genes gain and lose expression domains through a multi-step accumulation of small-scale mutations, such as point mutations or indels, that either create new or destroy old transcription factor binding sites (Stone and Wray, 2001). At the other extreme, a gene may evolve a regulatory element in a single step through the integration and cooption of a transposable element (TE) that harbors functional transcription factor binding sites (Bourque et al., 2008; El-Deiry et al., 1992; Jordan et al., 2003; Kunarso et al., 2010; Lynch et al., 2015; Polak and Domany, 2006; Wang et al., 2007). While there is ample data to support both models of gene regulatory evolution (reviewed in (Feschotte, 2008; Wray, 2007)), important questions with both models remain (Bourque et al., 2018). For example: Do TEs integrate with regulatory abilities, or as ‘pre-regulatory elements’ that need additional mutations to acquire regulatory functions? Do the functions of TE derived regulatory elements change, or is their evolution constrained by their functions upon integration? And does cooption of TEs into regulatory elements occur continuously or in a single wave? Answers to these (and other) questions are essential for understanding the contribution of TEs to gene regulatory evolution.

Extant mammals span several major evolutionary transitions during the origins, divergence, and conservation of pregnancy, including the origin of new cell-types in the uterine lining (endometrium), endometrial stromal fibroblasts (ESF), and decidual stromal cells (DSC) (Chavan et al., 2021; Erkenbrack et al., 2018; Wu et al., 2020), which mediate many of the maternal responses to pregnancy. For example, an essential step in the establishment and maintenance of pregnancy is the differentiation (decidualization) of ESFs into DSCs in response to progesterone acting through the progesterone receptor (PGR), the second messenger cyclic AMP (cAMP) acting through protein kinase A (PKA), the transcription factor FOXO1 (Gellersen and Brosens, 2003; Kajihara et al., 2013), and, in some species, to fetal signals (Gellersen et al., 2007). Decidualization evolved in the stem lineage of Eutherian mammals (Kin et al., 2015, 2014; Mess and Carter, 2006) and induces large-scale gene regulatory, cellular, and physiological changes in the endometrium that are essential for successful implantation and pregnancy in many eutherians, including humans.

We have previously shown that hundreds of genes gained and lost endometrial expression coincident with the origins of pregnancy and decidualization in early mammals (Lynch et al., 2015; Marinić et al., 2021), and that a second major episode of gene expression evolution occurred in primates (Mika et al., 2021a, 2021b). This latter wave of gene expression evolution occurred coincident with the origin of numerous primate-specific female reproductive and pregnancy traits including menstruation (Burley, 1979; Emera et al., 2012; Finn, 1998; Strassmann, 1996), decidualization in the absence of fetal signals (spontaneous decidualization) (Carter and Mess, 2017; Gellersen et al., 2007; Gellersen and Brosens, 2003; Kin et al., 2016, 2015; Mess and Carter, 2006), deeply invasive placentas (Carter et al., 2015; Pijnenborg et al., 2011a, 2011b; Soares et al., 2018), and a derived parturition signal (Csapo, 1956; Csapo and Pinto-Dantas, 1965). Remarkably TEs appear to have played an important role in the origins of pregnancy through cooption into progesterone-responsive cis-regulatory elements in DSCs and may have played a similar role during the evolution of primate-specific pregnancy traits, but we excluded these evolutionarily young TEs in our previous analyses (Lynch et al., 2015; Mika et al., 2021a). Here we expand on our previous studies on ancient mammalian TEs to all TEs and show that successive waves of TEs have been coopted into progesterone-responsive cis-regulatory elements, including a rolling wave of TE cooption in primates. Genes with regulatory elements derived from primate-specific TEs are among the most strongly differentially regulated by progesterone, have essential roles in decidualization, and likely contribute to primate-specific pregnancy traits such as spontaneous decidualization. Finally, we tested 89 consensus TE sequences, as a proxy for ancestral TE sequences, and found that nearly all have dominant repressor functions and latent enhancer functions in mammalian cells. These data suggest that primate-specific TEs played an important role in gene regulatory evolution in primate DSCs. Furthermore, our data suggest a general two-stage model of TE domestication into gene regulatory elements, whereby loss of ancestral repressor functions unmasks hidden enhancer functions.

## Results

### Transposable elements are major contributors to regulatory elements in DSCs

We have previously shown that Mammalian-, Therian-, and Eutherian-specific TEs played an important role in the origin of new *cis*-regulatory elements in DSCs during the evolution of pregnancy (Lynch et al., 2015)) and are enriched nearby genes that gained and lost expression in primate DSCs (Lynch et al., 2015; Mika et al., 2021a). Here we expanded these studies to all classes and ages of TEs using previously generated H3K4me3 ChIP-Seq, H3K27ac ChIP-Seq, FAIRE-Seq and DNaseI-Seq data to identify promoters, enhancers, and regions of open chromatin (Lynch et al., 2015; Mika et al., 2021a). We found that 58.7% of H3K27ac and 53.0% of H3K4me3 ChIP-Seq peaks, 42.2% of FAIRE-Seq peaks, and 67.2% of DNaseI-Seq peaks overlapped annotated transposable elements (**Figure 1A**). Next, we annotated these TEs by their lineage specificity and found that TEs from different age classes differentially contributed to each kind of regulatory element: relatively young (i.e., Primate-specific) TEs dominated the DNaseI, H3K27ac, and H3K4me3 datasets, whereas relatively ancient TEs (i.e., Eutherian-specific and older) were more common in the FAIRE dataset (**Figure 1B**). 427 TE families were enriched (eTE; >1.5-fold, *P*≤0.05, binomial test) within H3K27ac and H3K4me3 ChIP-Seq, and FAIRE-, DNase Seq peaks (**Figure 1C**), most of which were Eutherian- and Primate-specific (**Figure 1D**).

**Figure 1.**
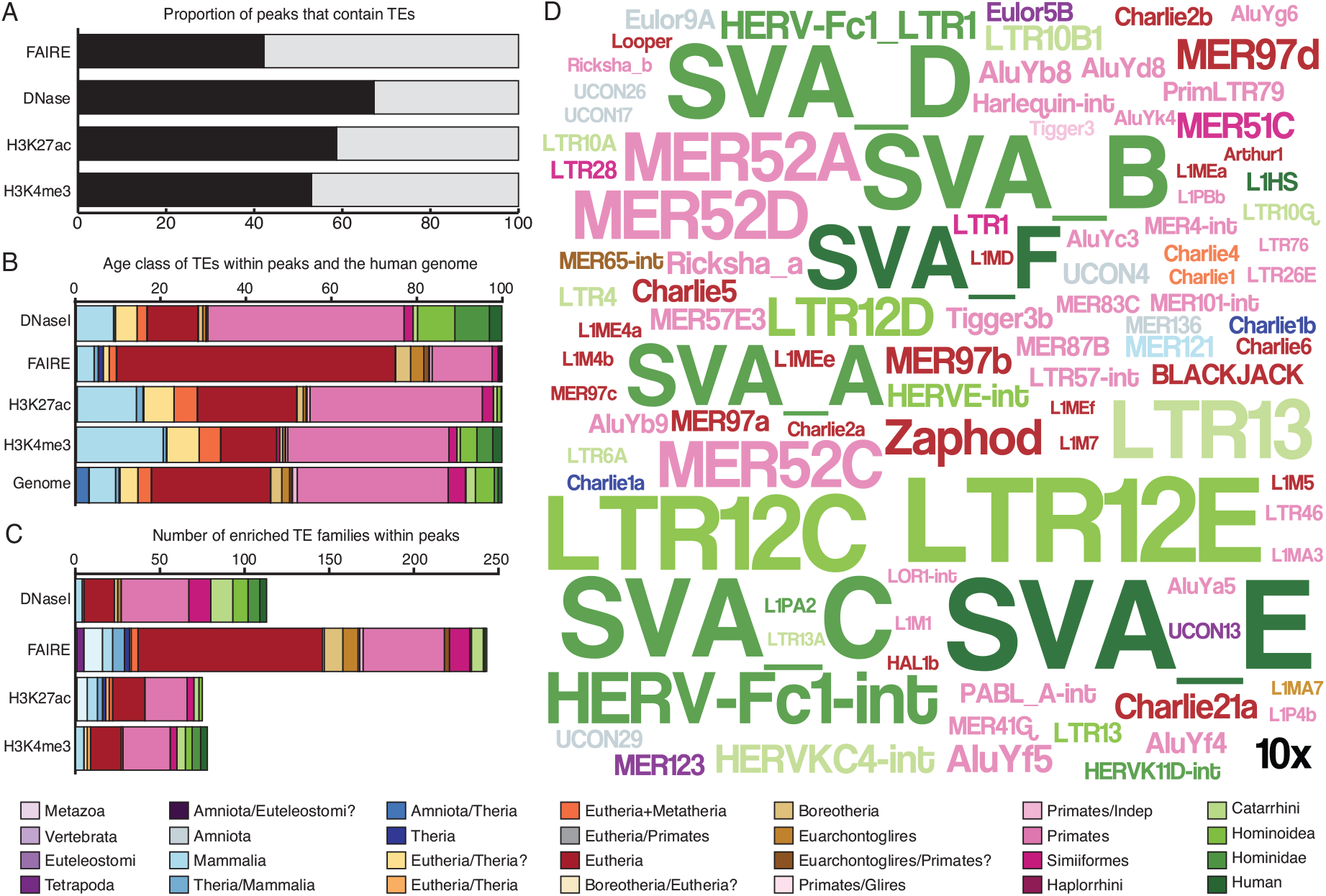
Transposable elements are major contributors to regulatory elements in endometrial stromal cells. (A) Proportion of DNaseI-Seq, FAIRE-Seq, H3K27ac ChIP-Seq, and H3K4me3 ChIP-Seq peaks that contain transposable elements (black) or no annotated transposable elements (gray). (B) Proportion of transposable elements of different ages (color coded by lineage specificity) within DNaseI-seq, FAIRE-seq, H3K27ac ChIP-seq, and H3K4me3 ChIP-Seq peaks and the human genome. (C) Number of transposable element families enriched within DNaseI-seq, FAIRE-seq, H3K27ac ChIP-seq, and H3K4me3 ChIP-Seq peaks compared to the human genome. (D) WordCloud of the 100 most enriched transposable elements within DNaseI-seq, FAIRE-seq, H3K27ac ChIP-seq, and H3K4me3 ChIP-Seq peaks. The size of the transposable element’s name corresponds to its enrichment (see inset 10-fold scale). Legend indicates the lineage-specificity (age) colored coding of transposable elements. **Figure 1 — Source data 1. Transposable elements enriched in DNaseI-Seq peaks in human DSCs.** **Figure 1 — Source data 2. Transposable elements enriched in FAIRE-Seq peaks in human DSCs.** **Figure 1 — Source data 3. Transposable elements enriched in H3K4me3 ChIP-Seq peaks in human DSCs.** **Figure 1 — Source data 4. Transposable elements enriched in H3K27ac ChIP-Seq peaks in human DSCs.** **Figure 1 — Source data 5. Summary enrichment data for the 427 eTEs.**

### TE-derived regulatory elements are enriched in transcription factor binding sites that regulates DSCs

To determine if TEs donated motifs for specific transcription factors, we identified over-represented transcription factor binding sites (TFBS) within eTE-derived regions of FAIRE-seq, DNase-seq, H3K27ac ChIP-Seq and H3K4me3 ChIP-Seq peaks using previously published ENCODE ChIP-Seq data for 132 transcription factors as well as previously published PGR ChIP-Seq data generated from human DSCs (Mazur et al., 2015). 53 TFBSs were enriched within regulatory eTEs relative to genomic TFBS abundances (FDR=0.05; **Figure 2A**), most notably PGR (enrichment=10.31, FDR<1.00×10^-250^), AHR (enrichment=2.00, FDR=1.00×10^-5^), and GATA (enrichment=1.25, FDR=1.30×10^-20^). We also observed enrichment for the KRAB-ZFPs ZNF263 (enrichment=1.71, FDR=7.20×10^-150^) and ZNF274 (enrichment=1.22, FDR=8.90×10^-3^), KAP1 (also known as TRIM28; enrichment=2.00, FDR=7.20×10^-41^), which binds KRAB-ZFPs and functions as a scaffold for the recruitment of histone modifying co-repressor complexes, and parts of the SWI/SNF chromatin remodeling complex such as BAF155 (enrichment=2.36, FDR=1.00×10^-76^), BAF170 (enrichment=1.93, FDR=4.20×10^-14^), INI1 (enrichment=1.43, FDR=9.10×10^-18^), and BRG1 (enrichment=1.38, FDR=2.70×10^-5^). These data suggest that TEs have donated binding sites for transcription factors that mediate decidualization such as PGR and its obligate co-factor GATA2 (Rubel et al., 2016, 2011), as well as general transcriptional repressors and chromatin modifying proteins (**Figure 2B**).

**Figure 2.**
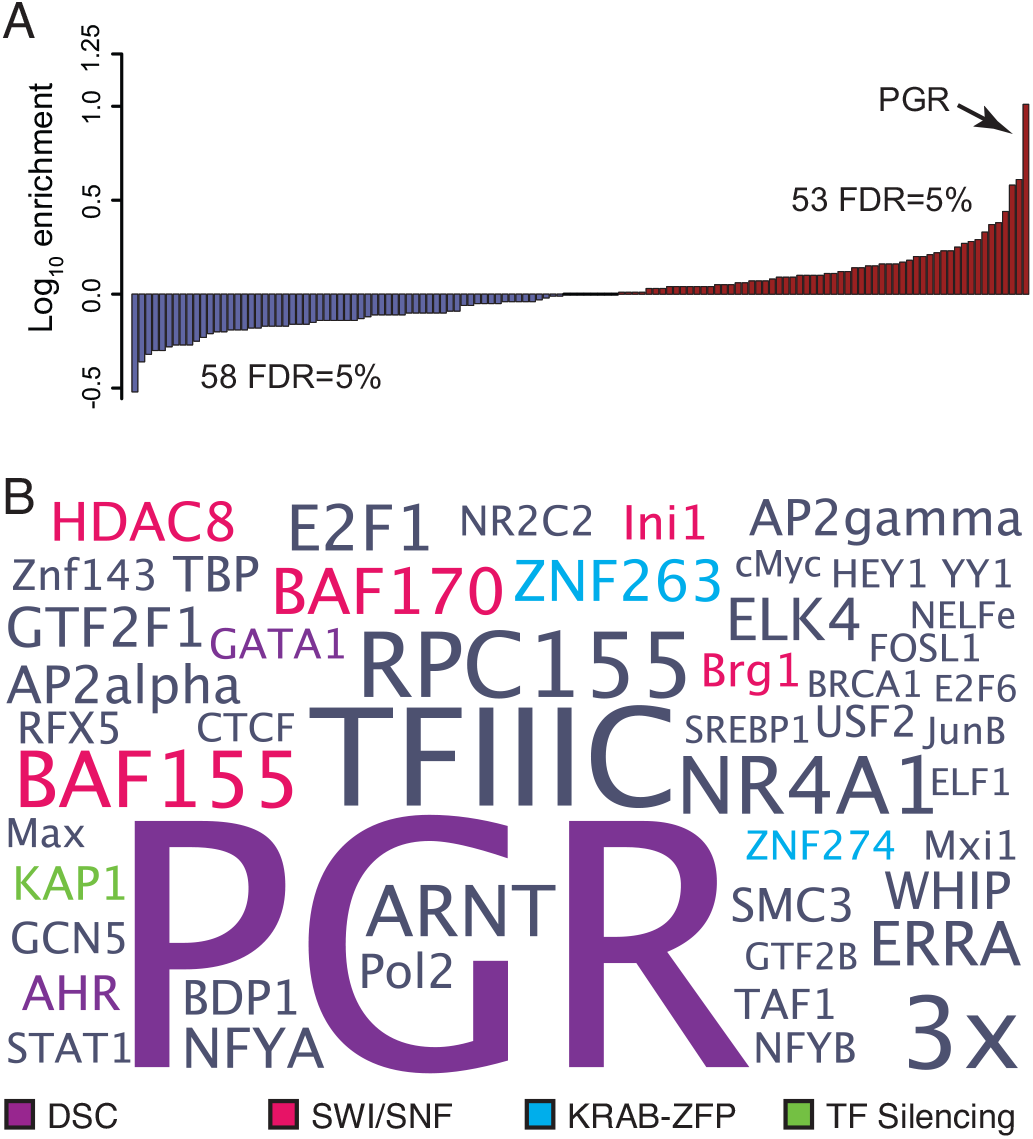
Transposable elements are enriched in binding sites for transcription factors that mediate hormone responsiveness and endometrial cell-type identity. (A) Distribution of enriched (red) and depleted (blue) transcription factor binding sites in transposable element derived segments of FAIRE-seq, DNaseI-seq, H3K27ac ChIP-seq, and H3K4me3 ChIP-Seq peaks relative to ChIP-Seq peaks with no transposable element overlap. 53 and 58 transcription factor binding sites were significantly enriched and depleted, respectively, at FDR=0.05%. (B) WordCloud of transcription factor binding site enrichment in transposable elements. Colors indicate transcription factors that mediate hormone responses (purple), remodel chromatin (pink), KRAB-ZFPs (blue), or involved in TE silencing (green). Data shown for ≥3-fold enriched transcription factors at FDR=0.05%. **Figure 2 — Source data 1. Transcription factors enriched in regulatory eTEs.**

### TE-derived regulatory elements augment ancient progesterone responsiveness

Our observation that specific TE families are enriched in DSC regulatory elements suggests that they may contribute to gene expression changes that occur during progesterone-induced decidualization and changes in decidualization-induced gene expression during human evolution. To test this hypothesis, we used parsimony to reconstruct the evolutionary history of gene expression in the pregnant uterus (**Figure 3A**) (Marinić et al., 2021), RNA-Seq data from human ESFs and DSCs to quantify gene expression changes induced by decidualization (**Figure 3B**), and previously published promoter capture HiC (pcHiC) data generated from human DSCs (Sakabe et al., 2020) to associate genes with putative regulatory elements (**Figure 3C**). Next, we used an F-test to compare the change in expression level (variance) between ESFs and DSCs between genes with and without TE-derived regulatory elements. We found that genes with eTE-derived regulatory elements were generally more strongly differentially regulated by decidualization than genes without eTE-derived regulatory elements (i.e., had greater expression variance), especially genes that were more recently recruited into endometrial expression (**Figure 3D**).

**Figure 3.**
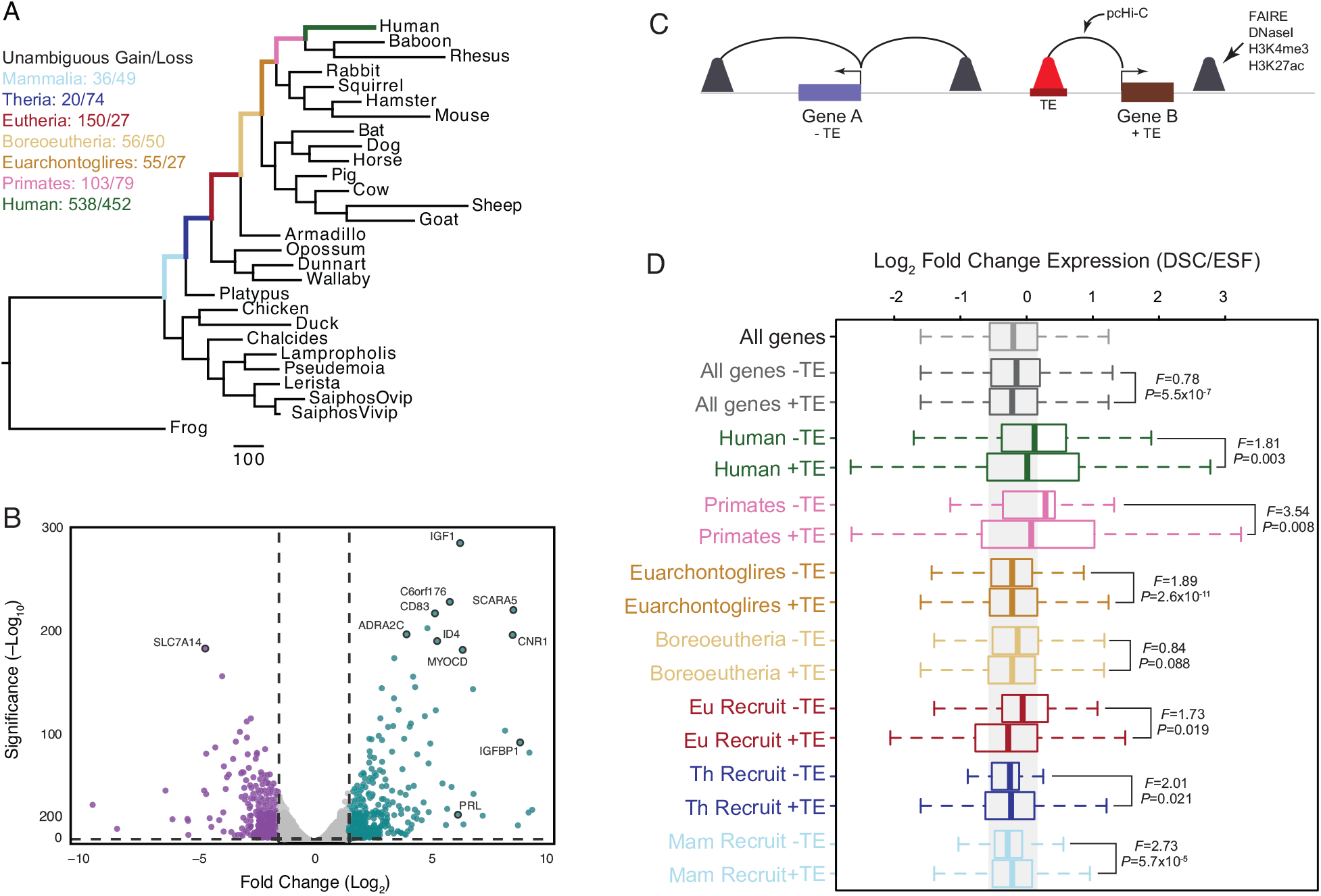
Genes associated with TE-derived regulatory elements are more strongly differentially regulated during decidualization than genes without TE-derived regulatory elements. (A) Parsimony reconstruction of gene expression gains and losses in the endometrium of amniotes. Numbers above branches indicate the average number of genes that gained and lost endometrial expression in that stem-lineage as inferred by parsimony. Branch lengths are drawn proportional to gene expression gain and loss events for all lineages. (B) Volcano chart showing genes differentially expressed between ESFs and DSCs. Upregulated genes are shown in blue, downregulated genes are shown in purple. Exemplar differentially expressed genes are indicated. (C) Cartoon of the pcHiC data to associate regulatory elements with nearby genes. (D) Recruited and ancestrally expressed genes associated with ancient mammalian TE-derived regulatory elements (+) are more strongly differentially regulated upon cAMP/MPA-induced decidualization than recruited or anciently expressed genes without TE-derived regulatory elements (-). F is the ratio of variances from a two-sample F-test. **Figure 3 — Source data 1. Parsimony reconstruction of genes that unambiguously gained and lost endometrial expression.** **Figure 3 — Source data 2. Gene expression changes induced by decidualization (from dataset GSE94036).** **Figure 3 — Source data 3. pcHiC data from human DSCs (from dataset SDY1626).**

### TE-derived regulatory elements are enriched in PGR binding sites

Our observations that TEs are enriched in PGR binding sites and are associated with genes that are strongly differentially expressed upon decidualization prompted us to explore the contribution of TEs to PGR binding sites in greater detail. We found that 62.8% (5344/8510) of PGR ChIP-Seq peaks in DSCs contained TEs (**Figure 4A**), nearly all of which are Mammalian-, Eutherian-, and Primate-specific (**Figure 4B**). PGR ChIP-Seq peaks, however, are almost exclusively enriched (>1.5-fold, *P*≤0.05, binomial test) in Eutherian- and Primate-specific TEs (**Figure 4C**). Consistent with a functional role for TE-derived PGR binding sites in orchestrating progesterone responsiveness, genes associated with TE-derived PGR binding sites by pcHiC (**Figure 4D**) were more strongly differentially expressed during decidualization than genes not associated with TE-derived PGR binding sites (**Figure 4E**); This trend was more pronounced for recently recruited genes (**Figure 4E**). We also found that genes associated by pcHiC with TE-derived PGR binding sites were significantly more dysregulated by siRNA-mediated PGR knockdown in DSCs (**Figure 4F**) than genes not associated with TE-derived PGR binding sites (**Figure 4G**).

**Figure 4.**
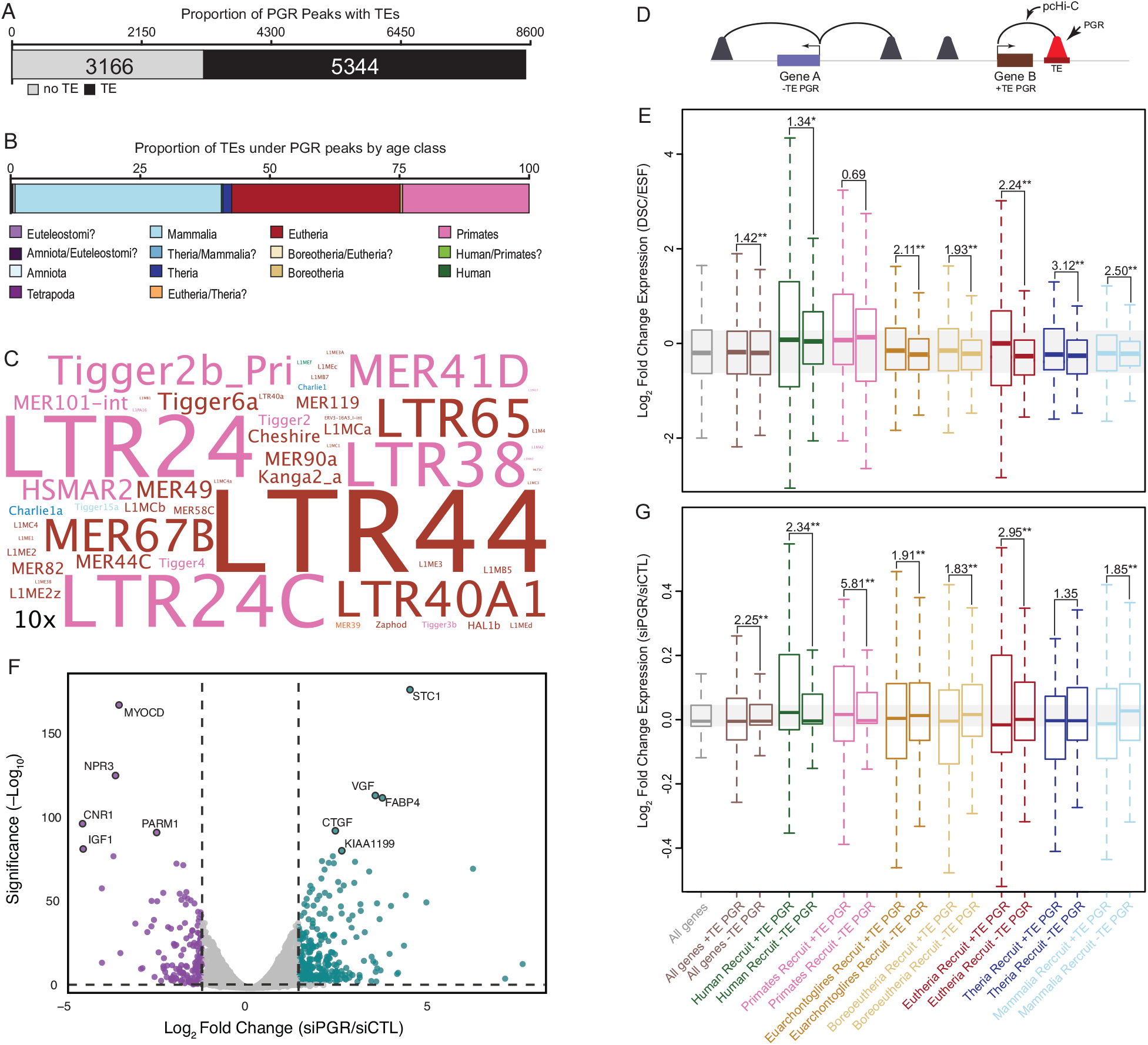
Lineage specific transposable elements remodeled progesterone receptor binding site architecture across the genome. (A) The number of PGR ChIP-Seq peaks that contain transposable elements in human decidual stromal cells. (B) Proportion of transposable elements in PGR ChIP-Seq peaks by age class. (C) WordCloud of TEs enriched in PGR ChIP-Seq peaks. Colors indicate age class. Inset scale (10x) shows 10-fold enrichment. (D) Cartoon of the pcHiC data to PGR binding sites with nearby genes. (E) Ancestrally expressed and recruited genes associated with TE-derived PGR binding sites (+ PGR) are more strongly differentially regulated by cAMP/MPA than genes without TE-derived PGR binding sites (-PGR). (F) Volcano chart showing genes differentially expressed between DSCs treated with control non-targeting siRNA and PGR-specific siRNA. Upregulated genes are shown in blue, downregulated genes are shown in purple. Exemplar differentially expressed genes are indicated. (G) Ancestrally expressed and recruited genes associated with TE-derived PGR binding sites (+ PGR) are more strongly dysregulated by PGR knockdown in human DSCs than genes not associated with ancient mammalian TE-derived PGR biding sites (-PGR). F is the ratio of variances from a two-sample F-test. Genes are grouped according to when they evolved endometrial expression. * = P<0.01, ** = P<1.0×10-5. **Figure 4 — Source data 1. Transposable elements enriched in PGR ChIP-Seq peaks in human DSCs.** **Figure 4 — Source data 2. Gene expression changes induced by siRNA mediated PGR knockdown (from dataset GSE94036).**

### TE-derived PGR binding sites in primates regulate genes essential for decidualization

To explore the functional consequences of eTE-derived PGR binding sites, we first binned eTEs into ancient mammalian or primate-specific categories based on our observation that TEs in regulatory elements are predominantly from two age classes. Next, we used the pcHiC data to associate genes with each category of eTE-derived PGR binding site and tested whether these genes were enriched in biology pathways (KEGG, Panther, Wikipathway, Reactome) using the over-representation analyses implemented in WebGestalt (Liao et al., 2019). Of the 965 genes associated with eTE-derived PGR binding sites expressed in DSCs (TPM≥2), 631 were associated with primate-specific eTE-derived PGR binding sites. These genes were enriched in 113 pathways at FDR≤0.10. Among the enriched pathways were many that play important roles in decidualization and pregnancy (**Table 1**), such as Wnt, FoxO, and prolactin signaling, and various pathways related to regulation of the cell cycle which plays a critical role in the earliest stages of decidualization (**Table 2**). In contrast, the 334 genes only associated with ancient mammalian eTE-derived PGR binding sites were not enriched in any pathway at at FDR≤0.10.

**Table 1.** Pathways in which genes regulated by primate-specific eTE-derived PGR bindingsites are enriched. E, enrichment ratio. P, hypergeometric P-value. FDR, Benjamini-Hochberg false discovery rate. Reference, reference for role of pathway in decidualization.

| Rank | Pathway | E | P | FDR | Reference |
| --- | --- | --- | --- | --- | --- |
| 2 | VEGFA-VEGFR2 Signaling Pathway | 2.60 | 2E-8 | 2.75E-5 | (Kim et al., 2013) |
| 6 | Insulin signaling pathway | 3.79 | 6E-7 | 2.72E-4 | (Kawamura et al., 2009) |
| 13 | TGF-beta Signaling Pathway | 3.52 | 6E-6 | 1.38E-3 | (Kim et al., 2005) |
| 17 | ErbB Signaling Pathway | 3.91 | 3E-5 | 4.28E-3 | (Klonisch et al., 2001) |
| 26 | Hippo signaling pathway | 3.02 | 5E-5 | 5.32E-3 | (Chen et al., 2017) |
| 27 | PDGF signaling pathway | 3.28 | 5E-5 | 5.41E-3 | (Schwenke et al., 2013) |
| 35 | FoxO signaling pathway | 3.11 | 9E-5 | 7.84E-3 | (Kajihara et al., 2013) |
| 37 | Mitophagy | 4.21 | 1E-4 | 8.71E-3 | (Mestre Citrinovitz et al., 2019) |
| 38 | Oncostatin M Signaling Pathway | 4.21 | 1E-4 | 8.71E-3 | (Fu et al., 2019) |
| 40 | TGF-beta Receptor Signaling | 4.56 | 1E-4 | 9.63E-3 | (Ni and Li, 2017) |
| 52 | BDNF signaling pathway | 2.85 | 3E-4 | 1.40E-2 | (Kawamura et al., 2009) |
| 54 | EGF/EGFR Signaling Pathway | 2.70 | 3E-4 | 1.54E-2 | (Large et al., 2014) |
| 55 | Wnt signaling pathway | 2.81 | 3E-4 | 1.54E-2 | (Zhang and Yan, 2016) |
| 65 | Prolactin Signaling Pathway | 3.60 | 4E-4 | 1.90E-2 | (Bao et al., 2007) |
| 74 | Myometrial Relaxation and Contraction Pathways | 2.63 | 6E-4 | 2.32E-2 | (Salomonis et al., 2005) |

**Table 2.** Pathways related to senescence and the cell-cycle in which genes regulated by primate-specific eTE-derived PGR binding-sites are enriched. E, enrichment ratio. P, hypergeometric P-value. FDR, Benjamini-Hochberg false discovery rate. Reference, reference for role of pathway in decidualization.

| Rank | Pathway | E | P | FDR | Reference |
| --- | --- | --- | --- | --- | --- |
| 11 | Cellular senescence | 3.25 | 6E-6 | 1.38E-3 | (Lucas et al., 2020) |
| 14 | DNA Damage Response (only ATM dependent) | 3.73 | 1E-5 | 2.23E-3 | (Lei et al., 2012) |
| 19 | G2/M Transition | 2.82 | 3E-5 | 4.37E-3 | (Logan et al., 2012) |
| 20 | Mitotic G2-G2/M phases | 2.79 | 3E-5 | 4.70E-3 | (Logan et al., 2012) |
| 41 | Apoptosis signaling pathway | 3.29 | 2E-4 | 1.08E-2 | (Joswig et al., 2003) |
| 42 | Oxidative Stress | 5.80 | 2E-4 | 1.08E-2 | (Yu et al., 2019) |
| 59 | Mitotic G1-G1/S phases | 2.75 | 4E-4 | 1.79E-2 | (Logan et al., 2012) |
| 66 | Cell Cycle | 2.96 | 4E-4 | 1.92E-2 | (Das, 2009) |
| 67 | Senescence and Autophagy in Cancer | 3.12 | 4E-4 | 1.92E-2 | (Deryabin et al., 2020) |
| 75 | Cell cycle | 2.87 | 6E-4 | 2.32E-2 | (Das, 2009) |
| 91 | Cyclin D associated events in G1 | 4.35 | 1E-3 | 3.09E-2 | (Logan et al., 2012) |
| 92 | G1 Phase | 4.35 | 1E-3 | 3.09E-2 | (Logan et al., 2012) |
| 99 | p53 signaling pathway | 3.42 | 1E-3 | 3.45E-2 | (Deng et al., 2016) |

### Consensus TEs are repressors with latent enhancer potential

It is not clear if TEs integrate with regulatory abilities, and therefore immediately function as regulatory elements, or if they integrate as ‘pre-regulatory elements’ that are weakly- or nonfunctional and require additional mutations to acquire regulatory functions. Alternatively, TEs might not initially have direct effects on the expression of nearby genes, rather they might have collateral effects on host gene transcription by being targets of TE silencing where a TE is bound by a host repressive transcription factor, thereby causing the silencing of nearby gene (Fueyo et al,. 2002). To test these scenarios, we selected 89 enriched TEs, synthesized their consensus sequences (conTE), and cloned them into the pGL3-Basic[minP] luciferase reporter vector. Next, we transiently transfected human ESFs and DSCs with each conTE reporter and used a dual luciferase reporter assay to test their regulatory abilities. We found that 55 (62%) the conTE reporters functioned as repressors in ESFs while 58 (65%) functioned as repressors in DSC (FDR≤0.05), in contrast only 13 (3%) and 21 (23%) had enhancer functions in ESFs and DSCs (**Figure 5A**; FDR ≤ 0.05). To test whether these effects were cell-type specific, we repeated the luciferase reporter assay in the human hepatocellular carcinoma cell line HepG2 and again observed that 59/89 (66%) were strong repressors (FDR≤0.05) whereas 28 were enhancers (**Figure 5A**; FDR ≤ 0.05). To determine if these results were species-specific we repeated the luciferase assay in mouse embryonic fibroblasts (MEFs) and observed that 51 (57%; FDR ≤ 0.05) of conTEs were repressors (34 were enhancers; FDR ≤ 0.05) whereas in elephant dermal fibroblasts 21 (24%; FDR ≤ 0.05) were repressors (34 were enhancers; FDR ≤ 0.05) (**Figure 5A**). While some conTEs, such as LTR elements, which have strong internal promoters, had enhancer functions in all cell-types, significantly more were repressors than expected by chance in ESFs (Binomial *P*=0.027), DSCs (Binomial *P*=1.00×10^-6^), HepG2 (Binomial *P*<1.00×10^-6^), and MEFs (Binomial *P*=0.027).

**Figure 5.**
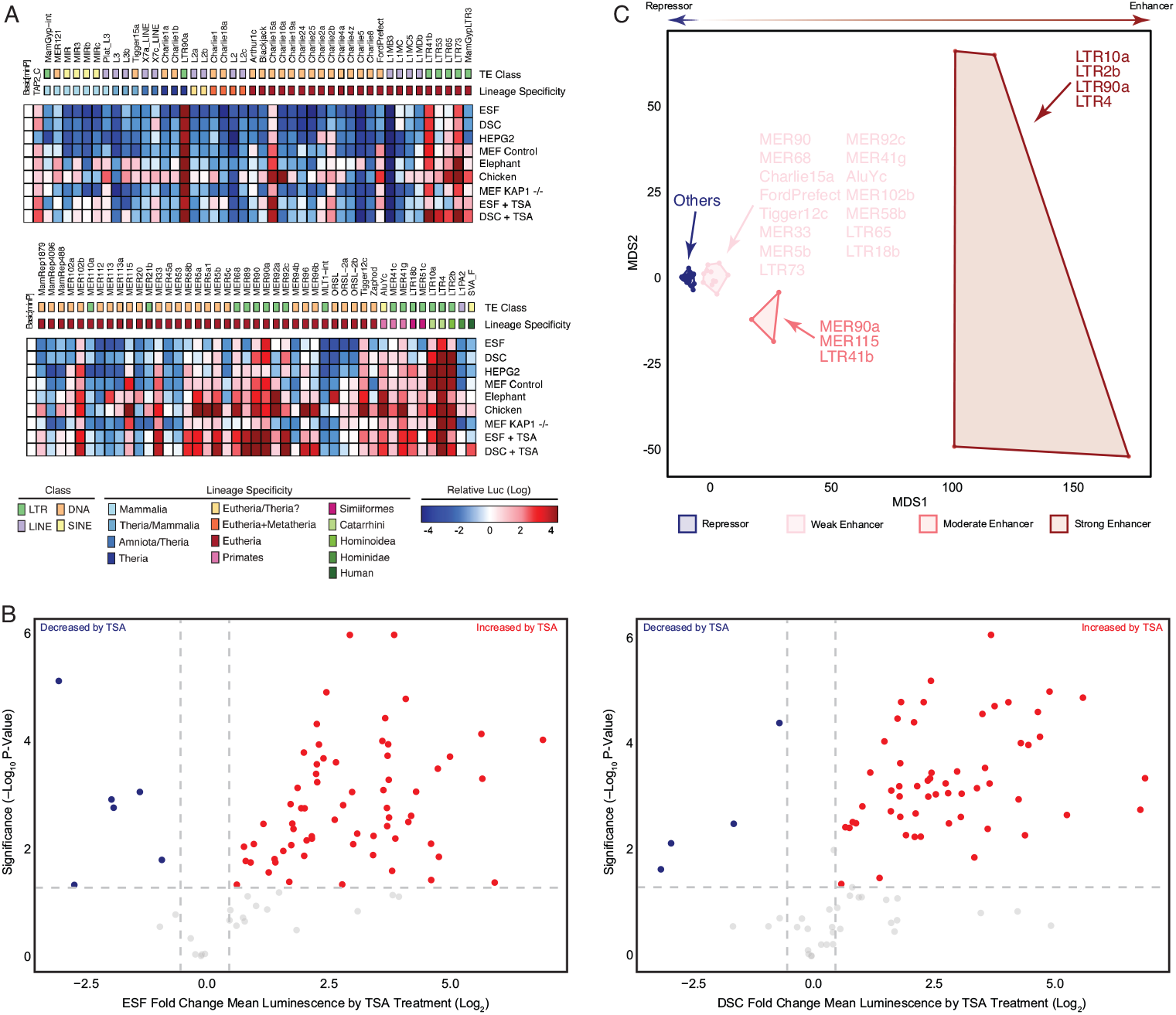
Consensus TEs are repressors with hidden enhancer potential. (A) Regulatory ability of consensus TEs was assessed using a dual luciferase reporter vector. Each box represents the mean luminescence values (log) of 4-6 replicate reporter assays. Basic[minP], empty reporter vector reference. TAP2_C is a known progesterone responsive enhancer^44^. The 89 TE constructs are sorted in order of decreasing age inferred by lineage specificity. Red indicates enhancer function and blue indicates repressor function (see inset color scale). ESF - Endometrial Stromal Fibroblasts, DSC - Decidual Stromal Cells, HEPG2 – Human Liver Cancer, MEF Control - Mouse Embryonic Fibroblast, MEF KAP1^-/-^ - Mouse *KAP1* knockout Embryonic Fibroblast, Elephant - Elephant Dermal Fibroblast, Chicken - Chicken Embryonic Fibroblast, +TSA - Trichostatin A added to media. (B) Volcano Plot ESF vs ESF treated with TSA and DSC vs DSC treated with TSA (C) Multidimensional scaling plot (MDS) of consensus TEs based on mean luminescence values (log) and grouped by K-means clustering (k=4). K-means clustering groups TEs into four categories corresponding to TEs with strong/moderate/weak enhancer or repressor functions.

Our observation that TEs enriched within regulatory elements and open chromatin are also enriched in binding sites for KRAB-ZFPs, KAP1, as well as histone modifying co-repressor complexes, suggest that conTEs may function as repressors because they are recognized by the host cell anti-TE machinery and silenced (Feschotte and Gilbert, 2012; Matsui et al., 2010; Rowe et al., 2010). To test this hypothesis, we repeated the luciferase reporter assay in ESFs and DSCs and treated cells with the mammalian class I and II histone deacetylase (HDAC) inhibitor trichostatin A (TSA). TSA treatment de-repressed 30/35 (86%; FDR ≤ 0.05) and 39/55 (71%; FDR ≤ 0.05) of the conTEs that were repressors in untreated ESFs and DSCs, respectively (**Figure 5B**). TSA treatment also unmasked the latent enhancer functions of 29 conTEs in ESFs and 26 conTEs in DSCs (for example: MIR3, MIRb, and MIRc, MamRep1879, and MER96b). Thus, while the majority of conTEs repress luciferase expression, their repressive abilities are likely HDAC dependent.

The preponderance of repressor functions among conTEs and their dependence on HDACs suggests that conTEs may be recognized by the KRAB-ZFPs/KAP1 TE suppression system and silenced by histone modifying co-repressor complexes (NuRD, CoREST, SWI/SNF) (Feschotte and Gilbert, 2012; Imbeault et al., 2017; Matsui et al., 2010; Rowe et al., 2010). To test this hypothesis, we repeated the luciferase reporter assay in *KAP1* knockout mouse embryonic fibroblasts (MEF *KAP1*^-/-^), but again observed 53 conTEs (59%; FDR ≤ 0.05) were repressors and relief of repression for only 8 (9%; FDR ≤ 0.05) conTEs (**Figure 5A**). To infer if repression may be mediated by KRAB-ZFPs, we took advantage of their restricted lineage specificity and tested the regulatory abilities of conTEs in chicken embryonic fibroblasts; The chicken genome only encodes 41 KRAB-ZFP, none of which are expressed in chicken embryonic fibroblasts (Addison et al., 2015). In stark contrast to the different mammalian cell-types we tested, only 18 (20%; FDR ≤ 0.05) of the conTEs functioned as repressors whereas 69 (78%; FDR ≤ 0.05) were strong enhancers in CEFs (**Figure 5A**).

Finally, we used multidimensional scaling to explore whether conTEs could be grouped into distinct clusters based on their regulatory abilities in different cell-types. We found that conTEs formed four clusters in the MDS plot, corresponding to those with weak, moderate, and strong enhancer functions and those with repressor functions (**Figure 5C**). Similarly, hierarchical clustering grouped conTEs with enhancer and repressor functions (**Figure 5 – figure supplement 1**). Consistent with our observation that the regulatory functions of conTEs were similar in different mammalian cell-types but different in CEFs, CEFs did not cluster with mammalian cell-types in an MDS plot (**Figure 5 – figure supplement 2**). Collectively these data indicate that while a few conTEs, mostly LTRs, have strong enhancer or promoter abilities, most conTEs function as HDAC-dependent repressors in mammalian cells. We note that these effects were independent of conTE length across all experiments indicating there is no bias for short or long TEs to act as repressors or enhancers of luciferase expression (**Figure 5 – Figure supplement 1**).

Our observations that TEs are enriched binding sites for transcription factors important for DSC function and that consensus TEs have dominant repressor abilities and latent enhancer potential, suggest that these consensus TEs should have binding sites for repressors and activators. To explore this possibility, we used XSTREME to discover over-represented motifs in conTEs. Consistent with the repressor abilities of conTEs, we found that zinc finger proteins were among the most numerous over-represented motifs, including matches binding sites for KAP1-interacting zinc finger proteins such as ZNF320, ZIC1-3, ZNF417, ZKSCAN5, and ZNF189, as well as repressors and co-repressors such as CTCF, EHF, ESRRA, ESRRB, and ESRRG (**Figure 5 – Figure supplement 4-6**). We also found over-represented motifs that match binding sites for transcription factors important for regulating stromal cell identity and progesterone responsiveness, such as PGR, FOXO, and Hox. These data suggest that the ability of conTEs to act as repressors is dependent on pre-existing binding sites for transcription factors with repressor functions, either directly like ESRRA-G, indirectly though recruiting TE silencing repressors mediated by KRAB-ZFP containing proteins, or indirectly through regulating chromatin organization and positional information mediated by CTCF.

## Discussion

### Relationship to previous studies

We previously used a combination of comparative transcriptomics and genomics to show that ancient mammalian transposable elements, i.e., TEs that integrated into the genome of the last common ancestor of Mammalia, Theria, and Eutheria, were enriched within active regulatory elements and progesterone receptor binding sites in human decidual stromal cells (Lynch et al., 2015). However, in that study we focused exclusively on ancient mammalian TEs and used a computational method (GREAT), which assigns each gene a regulatory domain including a basal domain that extends 5 kb upstream and 1 kb downstream from its transcription start site and an extension up to the basal regulatory domain of the nearest upstream and downstream genes within 1 Mb, to associate TE-derived regulatory elements with genes. Here we considered all TE age classes, rather than just ancient mammalian ones, and use a direct experimental method, promoter capture HiC (pcHiC), to directly associate TE-derived regulatory elements with genes expressed in human decidual stromal cells. This more expansive TE inclusion set and the direct association of genes with their long-range regulatory elements suggests transposable elements continuously remodel the regulatory landscape, transcriptome, and function of decidual stromal cells.

### Two waves of transposable element cooption remodel the transcriptome and regulatory landscape of decidual stromal cells

Transposable elements are so frequently coopted into regulatory elements that it is not possible to cite all or even most studies reporting either the cooption of individual TEs or large-scale cooption of (almost) entire TE families. Previous studies, for example, have dissected in great detail the cis-regulatory element that drives extra-pituitary *prolactin* (*PRL*) in human decidual stromal cells (DSCs), which is normally expressed by the pituitary and immune cell-types (Gerlo et al., 2006). These studies found that human *PRL* expression is initiated from an alternative promoter located 5.8kb upstream of the canonical pituitary transcription start site which contains binding sites for transcription factors that are essential for the identity and function of DSCs including PGR, FOXO1A, ETS1, CEBPB, and FOS (Gerlo et al., 2006). Remarkably, the human decidual *PRL* promoter is derived from a primate-specific long terminal repeat (LTR)-like transposable element in the medium reiterated repeat (MER39) family and an upstream enhancer derived from a Eutherian-specific MER1 class of DNA transposon (MER20) (Gerlo et al., 2006). However, while there is little evidence that other MER39 elements function as promoters (Emera and Wagner, 2012), MER20 elements function as enhancers in DSCs for numerous genes across the genome (Lynch et al., 2011).

These data suggest that Eutherian-specific transposable elements played a role in re-wiring the gene regulatory network during the evolution of pregnancy and decidualization. Consistent with this observation, we previously found large-scale cooption of Mammalian-, Therian-, and Eutherian-specific TEs (AncMamTEs) into progesterone responsive cis-regulatory elements. Here we expanded on these studies and found that 427 TE families were enriched in DSC regulatory elements, nearly half of which were primate-specific. Thus, there were at least two waves of TE cooption into decidual regulatory elements – a first wave in early mammals and a second wave in primates. Remarkably, genes associated by pcHiC with TE-derived regulatory elements were significantly more responsive to progesterone than genes without TE derived cis-regulatory elements. Furthermore, genes with primate-specific TE-derived PGR binding sites were more progesterone responsive than genes with either ancient TE-derived PGR binding sites or genes without TE-derived PGR binding sites. These data suggest that TEs may have played a role in rewiring the progesterone responsive gene regulatory network both during the evolution of pregnancy in early mammals and in primates.

### Primate-specific transposable elements contribute to the function of decidual stromal cells

While eutherian mammals share a suite of traits mediated by endometrial stromal lineage cells that support prolonged pregnancies, there is also considerable variation in pregnancy traits within eutherians. Catarrhine primates, for example, have evolved spontaneous decidualization (differentiation) of endometrial stromal fibroblasts (ESFs) into decidual stromal cells (DSCs) under the combined action of progesterone, cyclic adenosine monophosphate (cAMP), and other unknown maternal signals (Carter and Mess, 2017; Gellersen et al., 2007; Gellersen and Brosens, 2003; Kin et al., 2016, 2015; Mess and Carter, 2006). Decidualization induces dramatic gene expression and functional changes (Aghajanova et al., 2011; Gellersen et al., 2007; Giudice, 2003), but the molecular mechanisms that underlie the evolution of spontaneous decidualization are largely unknown. We found that coopted primate-specific TEs regulate genes in several pathways involved in decidualization and the cell cycle (**Table 1** and **Table 2**), most notably the FOXO1 signaling pathway. For example, the transcription factor FOXO1 plays a particularly important function as a key regulator of decidualization, which integrates cAMP and progesterone signaling through physical and functional interactions with the progesterone receptor and other transcription factors to direct expression of progesterone response genes (Gellersen and Brosens, 2003; Lynch et al., 2009; Takano et al., 2007). Similarly, among the earliest steps in the decidualization process is cell cycle exit (Das, 2009; Logan et al., 2010); progesterone initially induces cell cycle arrest at the G0/G1 checkpoint followed by arrest at the G2/M checkpoint, both of which are regulated by p53 signaling (Logan et al., 2012). These data suggest that primate-specific TEs may have played a role in the origin of spontaneous decidualization by altering the regulation of genes in the FOXO1 signaling and cell cycle regulation pathways.

### A multi-stage model for transposable element domestication

Numerous studies have shown that transposable elements have donated binding sites for transcription factors to the genome, can be bound by transcription factors, and have been coopted into cis-regulatory elements (El-Deiry et al., 1992; Jordan et al., 2003; Kunarso et al., 2010; Lynch et al., 2011; Polak and Domany, 2006; Wang et al., 2007), but it generally has not been determined if TEs integrate into the genome with regulatory functions and therefore immediately function as regulatory elements or if they integrate as ‘pre-regulatory elements’ that are not immediately functional and require additional mutations to acquire regulatory functions. We addressed this question using luciferase assay functional tests of conTEs and found the majority of consensus TEs have enhancer abilities but these abilities are silenced in mammalian cells, perhaps by KRAB-ZFPs and/or the NURD HDAC inhibitory complex. We do see a mild enrichment within the dataset of TEs with regulatory marks in DSCs. This enrichment is likely low because 1) the elements analyzed have likely escaped repression by KRAB-ZFPs in order to be coopted into enhancers, and 2) the datasets used to identify KRAB-ZFP, KAP1, and the majority of other transcription factors, were from ENCODE and not DSCs. However, the relief of repression seen in chicken cells, which do not express KRAB-ZFPs (Addison et al., 2015), is at least coincidental evidence that KRAB-ZFPs may play a role in silencing conTEs. In embryonic stem cells, KAP1 binds the KRAB-ZFPs and coordinates the silencing of the bound TE (Feschotte and Gilbert, 2012; Matsui et al., 2010; Rowe et al., 2010). However, in adult somatic tissues the role of KAP1 is unclear and TE silencing in these tissues may be independent of KAP1 but require other housekeeping co-repressors (Ecco et al., 2016; Matsui et al., 2010; Rowe et al., 2010); In this model KRAB-ZFPs bind TEs in somatic cells, but silence TEs by recruiting repressors other than KAP1. Thus our observations suggest that silencing of the conTEs in MEFs may require KRAB-ZFPs but be KAP1 independent. These results need to be confirmed, for example, by ChIP to demonstrate conTEs are indeed bound by KRAB-ZFPs. We hypothesize that only when TEs escape this silencing regulation are they coopted by the genome to play a regulatory role (**Figure 6**).

**Figure 6.**
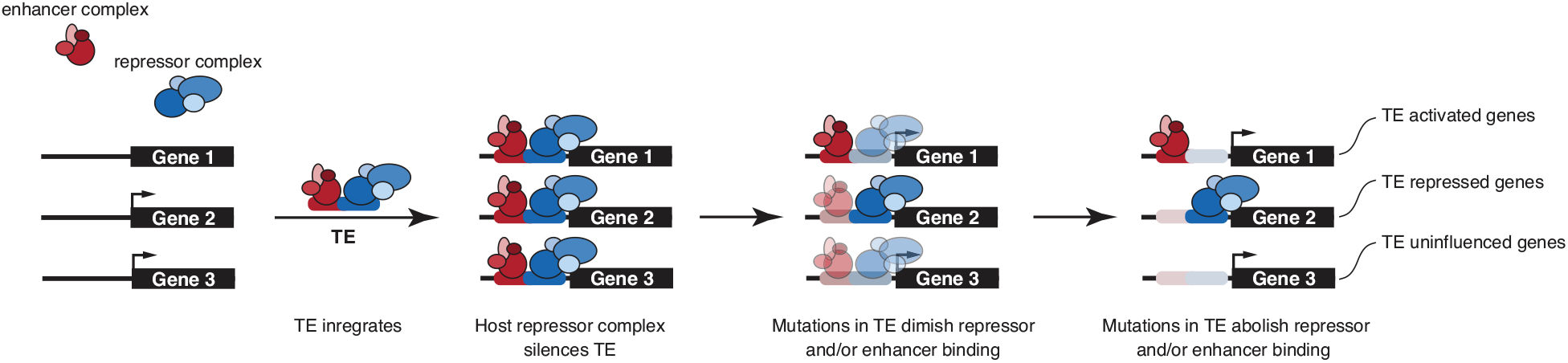
A multi-stage model for transposable element domestication. In this model, a gene may (arrowhead) or may not be expressed in a cell-type. Upon integration into the host genome, a TE brings with it potential transcription factor binding sites to assemble an enhancer complex, but this innate enhancer ability is blocked by host repressor complexes that generally silence the transposition of TEs. After the active transposition phase of the TE lifespan has ended, selection to maintain binding sites for the repressor complex is lost unmasking latent enhancer functions. Alternatively, TEs may maintain binding sites for repressors to acts as repressor elements or may lost both enhancer and repressor binding sites and not influence gene expression.

### Caveats and limitations

Ideally, we would use an ancestral sequence reconstruction (ASR) of each TE to determine if TEs had ancestral regulatory abilities. However, reconstructing the earliest ancestral sequence for most TEs is not possible because most TEs do not have an outgroup to root their phylogeny and therefore we cannot identify which node is the deepest ancestor. In place of an ASR, we and others who have explored similar questions used conTEs. This introduces an obvious limitation to our inferences: the conTE may not be an accurate representation of the deepest ancestral sequence. Indeed, there is no guarantee that a consensus sequence represents a sequence that ever existed. Another possible limitation of our approach is the use of immortalized ESFs maintained under standard tissue conditions for both functional genomic studies and luciferase assays, which may not faithfully represent *in vivo* functions. These limitations impact virtually every study of primate pregnancy, however, endometrial organoids and iPSC-derived endometrial stromal fibroblasts are promising systems in which to study the evolution of pregnancy (Abbas et al., 2020; Boretto et al., 2017; Marinić et al., 2020; Rawlings et al., 2021; Turco et al., 2017). Another limitation of our experimental approach is that the reporter assays used an episomal vector rather than a construct inserted into the genome which may bias our results by removing context dependent effects of regulatory elements within the genome and it is possible that TSA treatment induces the activation of a general transcriptional activator leading to up-regulation of luciferase expression upon TSA treatment. Finally, we have not performed the kinds of empirical studies necessary to confirm a causal relationship between TEs and gene expression levels, however, our results provide strong circumstantial evidence that TEs have influenced the gene expression and function of decidual cells.

## Conclusions

Here we demonstrate that there were two major waves of TE cooption into regulatory elements in endometrial stromal lineage cells, specifically in ancient mammals and primates, and that these TEs donated functional transcription factor binding sites to the genome. Furthermore, the majority of the TEs tested have context-dependent repressor and enhancer functions, suggesting they may have integrated into the genome with regulatory abilities and that their genomic functions are cell-type dependent. Genes regulated by TE-derived regulatory elements are among the most progesterone responsive in the genome and are associated with essential functions of stromal cells and the processes of decidualization. These data suggest that TEs have played an important role in the evolution of gene regulation and function of endometrial stromal lineage cells. This may have had a particularly significant effect in primates, which have divergent pregnancy traits than other mammals.

## Materials and Methods

### Identification of TE containing regulatory elements

We used previously published ChIP-Seq data generated from human DSCs that were downloaded from NCBI SRA and processed remotely using Galaxy (Afgan et al., 2016). ChIP-Seq reads were mapped to the human genome (GRCh37/hg19) using HISAT2 (Kim et al., 2019, 2015; Pertea et al., 2016) with default parameters and peaks called with MACS2 (Feng et al., 2012; Zhang et al., 2008) with default parameters. Samples included H3K4me3 (GSE61793), H3K27ac (GSE61793), PGR (GSE69539), the PGR A and B isoforms (GSE62475), and DNase1 - Seq (GSE61793). FAIRE-Seq peaks were downloaded from the UCSC genome browser and not re-called.

To identify regulatory elements derived from transposable elements (TEs), peaks were intersected with the RepMask 3.2.7 track at the UCSC genome browser (repbase libraries release 20050112). Non-transposable element annotations were removed and corrected for fragmented annotations. To identify TEs that were significantly enriched within TE-derived peaks the TEanalysis pipeline (https://github.com/4ureliek/TEanalysis) was used with 10,000 replicates. A custom bioinformatic pipeline was used to determine enrichment of transcription factor binding sites in TEs that intersect with the DSC FAIRE-seq, DNase-seq, H3K27ac ChIP-Seq, and H3K4me3, and PGR ChIP-Seq peaks (https://github.com/4ureliek/TEanalysis) (Lynch et al., 2015) versus the genomic abundance of ChIP-Seq peaks; 10,000 bootstrap reshufflings were used to assess statistical significance. Scripts are publicly available and archived at https://github.com/4ureliek/TEanalysis. The location of ChIP-Seq peaks in hg19 ENCODE data was downloaded from the USCS genome browser (Txn Fac ChIP V2 - Transcription Factor ChIP-Seq from ENCODE (V2)). The location of PGR ChIP-Seq peaks was obtained from GEO (GSE94036) and is available (Mazur et al., 2015). We also used previously published promoter capture HiC (pcHiC) generated from DSCs (Sakabe et al., 2020) to associate genes with regulatory elements (SDY1626) and did not reanalyze these data.

### Gene expression data and parsimony reconstruction of gene expression gain/loss

We used previously generated RNA-Seq data from the pregnant or gravid uterus of amniotes (**Supplementary Table 1**) and Kallisto version 0.42.4 (Bray et al., 2016) to quantify gene expression levels. Kallisto was run with default parameters, bias correction, 100 bootstrap replicates. Kallisto outputs are gene expression levels in transcripts per million (TPM). Our previous studies of endometrial gene expression data suggests that genes with a TPM≤2 are likely from transcriptionally suppressed genes (Wagner et al., 2013). This threshold is consistent with one obtained by comparing the transcript abundance with the chromatin state of the respective gene (Hebenstreit et al., 2011) and classified genes with TPM ≥ 2 as expressed and those with TPM<2 as not expressed. Next, we used Mesquite (v2.75) and parsimony optimization to reconstruct ancestral gene expression states and identified genes that gained or lost endometrial expression in Amniotes. Expression was classified as an unambiguous gain if a gene was not inferred as expressed at the ancestral node (state 0) but inferred as expressed in a descendent node (state 1) and vice versa for the classification of a loss from endometrial expression.

### Gene expression data from human ESFs and DSCs

We used previously published RNA-Seq gene expression data generated from human primary human ESFs treated for 48 hr with control non-targeting and PGR-targeting siRNA prior to decidualization stimulus for 72 hr (GSE94036) to compare treatments (ESF/DSC and DSC+control siRNA/DSC+PGR siRNA) and gene sets (+TE/-TE). Data were downloaded from NCBI SRA and processed remotely using the Galaxy platform (https://usegalaxy.org/; Version 20.01). RNA-Seq datasets were transferred from SRA to Galaxy using the Download and Extract Reads in FASTA/Q format from NCBI SRA tool (version 2.10.4+galaxy1). We used HISAT2 (version 2.1.0+galaxy5) to align reads to the Human hg38 reference genome using single- or paired-end options depending on the dataset and unstranded reads, and report alignments tailored for transcript assemblers including StringTie. Transcripts were assembled and quantified using StringTie (v1.3.6) (Pertea et al., 2016, 2015), with reference file to guide assembly and the “reference transcripts only” option, and output count files for differential expression with DESeq2/edgeR/limma-voom. Differentially expressed genes were identified using DESeq2 (Love et al., 2014) (version 2.11.40.6+galaxy1). The reference file for StringTie guided assembly was wgEncodeGencodeBasicV33. Comparisons between Log2 fold change in gene expression between ESFs and DSCs were estimated by identifying genes with and without TE derived regulatory elements and comparing the variance in fold change expression between these two sets with an F-test in R.

### Over Representation Analyses (ORA)

We used WebGestalt v. 2019 (Liao et al., 2019) to identify enriched ontology terms using over-representation analysis (ORA). We used ORA to identify enriched terms for three pathway databases (KEGG, Reactome, and Wikipathway), three disease databases (Disgenet, OMIM, and GLAD4U), and a custom database of genes implicated in preterm birth by GWAS. The preterm birth gene set was assembled from the NHGRI-EBI Catalog of published genome-wide association studies (GWAS Catalog), including genes implicated in GWAS with either the ontology terms “Preterm Birth” (EFO_0003917) or “Spontaneous Preterm Birth” (EFO_0006917), as well as two recent preterm birth GWAS (Sakabe et al., 2020; Warrington et al., 2019) using a genome-wide significant P-value of 9×10-6. The custom gmt file used to test for enrichment of preterm birth associated genes is included as a supplementary data file to Figure 2 (Figure 2 — Source data 1).

### Transposable element reporter vectors

To generate luciferase reporter vectors for functional testing, we selected 79 of the 427 TE families enriched within DSC regulatory regions as marked H3K4me3 ChIP-seq, H3K27ac ChIP-seq, DNaseI-seq, and FAIRE-Seq datasets. These 79 TEs also met the following criteria: 1) At least 1 element from every lineage; and 2) represented all 4 classes of TEs (LTR, LINE, SINE, and DNA). Consensus sequences for these elements were taken from the database Dfam (Wheeler et al., 2013). 10 additional TEs also found in the enriched 427 were then chosen that were unique to Old World Monkeys or younger and their consensus sequences were also obtained from Dfam (Wheeler et al., 2013). The total set of elements were biased towards, but not limited to, the DNA class of transposable elements. These 89 consensus sequences were then synthesized by Genscript and cloned into the pGL3 Basic vector (Promega) with an added minimal promoter (pGL3Basic[minP]) and are available in the supplementary materials.

### Cell lines

Human hTERT-immortalized endometrial stromal fibroblasts were purchased from ATCC (CRL-4003), their identity has been authenticated by ATCC and were determined by the Lynch lab to be mycoplasma free. *KAP1* knockout mouse embryonic fibroblasts (MEFs) and Flox/Flox control MEFs were gift from D. Trono (Ecole Polytechnique Fédérale de Lausanne), their identity has been authenticated by the Trono lab and were determined by the Lynch lab to be mycoplasma free. Chicken embryonic fibroblast cells were purchased from ATCC (CRL-12203), their identity has been authenticated by ATCC and were determined by the Lynch lab to be mycoplasma free. HEPG2 cells were a gift from C. Brown (University of Pennsylvania), their identity has been authenticated by the Brown lab and were determined by the Lynch lab to be mycoplasma free. African elephant fibroblasts were a gift from the San Diego Frozen Zoo, their identity has been authenticated by the San Diego Frozen Zoo and were determined by the Lynch lab to be mycoplasma free.

### Cell culture and luciferase assays

Endometrial stromal fibroblasts, *KAP1* knockout mouse embryonic fibroblasts (MEFs), and Flox/Flox control MEFs were maintained in phenol red free DMEM (Gibco) supplemented with 10% charcoal stripped fetal bovine serum (CSFBS; Gibco), 1x ITS (Gibco), 1% sodium pyruvate (Gibco), and 1% L-glutamine (Gibco). Chicken embryonic fibroblast cells (ATCC CRL-12203) and HEPG2 cells were maintained in phenol red containing DMEM + Glutamax (Gibco) supplemented with 10% fetal bovine serum (FBS, Gibco) and Normocin (InviviGen). Elephant fibroblasts were maintained in 1:1 MEM (Corning cellgro) supplemented with 10% FBS, 1% penstrep (Gibco), 1x sodium pyruvate (Gibco), and 1x L-glutamine (Gibco) to FGM-2 (Lonza), made per manufacturer’s instructions.

Confluent cells in 96 well plates in 80μl of Opti-MEM (Gibco) were transfected with 100ng of the TE containing luciferase plasmid and 10ng of the pRL-null renilla vector (Promega) with 0.1μl PLUS reagent (Invitrogen) and 0.25μl of Lipofectamine LTX (Invitrogen) in 20μl Opti-MEM. The cells incubated in the transfection mixture for 6hrs and the media was replaced with the maintenance media overnight. Decidualization of ESFs was then induced by incubating the cells in the decidualization media: DMEM with phenol red (Gibco), 2% CSFBS (Gibco), 1% sodium pyruvate (Gibco), 0.5mM 8-Br-cAMP (Sigma), and 1μM of the progesterone analog medroxyprogesterone acetate (Sigma) for 48hrs. ESFs (decidualization controls) were incubated in the decidualization control media (phenol red free DMEM (Gibco), 2% CSFBS (Gibco), and 1% sodium pyruvate (Gibco) instead for 48hrs. For trichostatin A (TSA; Tocris Bioscience) trials, 1μM TSA was added to all the medias from plating through decidualization. After decidualization for ESFs and DSCs or after 48hrs from transfection for other cell types, Dual Luciferase Reporter Assays (Promega) were started by incubating the cells for 15mins in 20μl of 1x passive lysis buffer. Luciferase and renilla activity were then measured using the Glomax multi+ detection system (Promega). Luciferase activity values were standardized by the renilla activity values and background activity values as determined by measuring luminescence from the pGL3-Basic[minP] plasmid with no insert. Each luciferase experiment was replicated in 4-6 independent experiments. To identify significant expression shifts, we performed Wilcoxon tests on the data, and adjusted by the Benjamini and Hochberg (Benjamini and Hochberg, 1995) method for multiple testing; significance was determined by having an adjusted p-value ≤ 0.05.

### Data exploration and Multi-Dimensional Scaling (MDS)

We used classical Multi-Dimensional Scaling (MDS) to explore the structure of luciferase assay data. MDS is a multivariate data analysis method that can be used to visualize the similarity/dissimilarity between samples by plotting data points onto two-dimensional plots. MDS returns an optimal solution that represents the data in a two-dimensional space, with the number of dimensions (k) specified a priori. Classical MDS preserves the original distance metric, between data points, as well as possible. MDS was performed using the vegan R package (Oksanen et al., 2019) with four reduced dimensions. Luciferase assay data were grouped using K-means clustering with K=2-6, K=4 optimized the number of distinct clusters and cluster memberships.

### Motif discovery in conTEs

We used XSTREME (Motif Discovery and Enrichment Analysis) version 5.5.0, implemented in the MEME suite of tools (https://meme-suite.org/meme/) to identify sequence motifs that were over-represented in conTE sequences using three runs: 1) Default settings; 2) Default settings relaxed to allow for up to 15 motifs; and 3) To specifically search for binding sites similar to PGR, we note that the PGR binding site is identical to the NR3C2 binding site. Discovered motifs were matched to known motifs in the JASPAR 2022 database. Exact search parameters are given in the output files **Figure 5 – Figure supplement 4-6.**

## Author Contributions

KM and VJL collaboratively conceived, designed, interpreted data, and wrote this paper, as well as analyzed the expression shifts of genes associated with TE derived regulators or TE derived PGR binding sites in response to progesterone signaling and did the binding site enrichment analyses. KM analyzed the RNA-Seq data, performed the gene expression calling, completed the parsimony reconstruction, and prepped, conducted, and analyzed the luciferase assays and associated cell culture. VJL identified the TE containing regulatory elements.

## Acknowledgements

The authors thank A. Kapusta for assistance with running the TE enrichment program, D. Trono (Ecole Polytechnique Fédérale de Lausanne) for providing the *KAP* KO and control MEFs, C. Brown (University of Pennsylvania) for providing the HEPG2 cells, and N. Sakabe (University at Chicago) for providing the pcHiC interaction data. This study was supported by a grant from the March of Dimes (March of Dimes Prematurity Research Center to principal investigator VJL), a Burroughs Welcome Fund Preterm Birth Initiative grant (1013760, to principal investigator VJL), and a Burroughs Welcome Fund Next Gen Pregnancy Research Grant (1022513, to principal investigator VJL). The funders had no role in study design, data collection and analysis, decision to publish, or preparation of the manuscript.

## Data availability statement

All data necessary to reproduce the results reported in this study are publicly available or available in the supplementary materials.

---

**Figure 5 – figure supplement 1.**
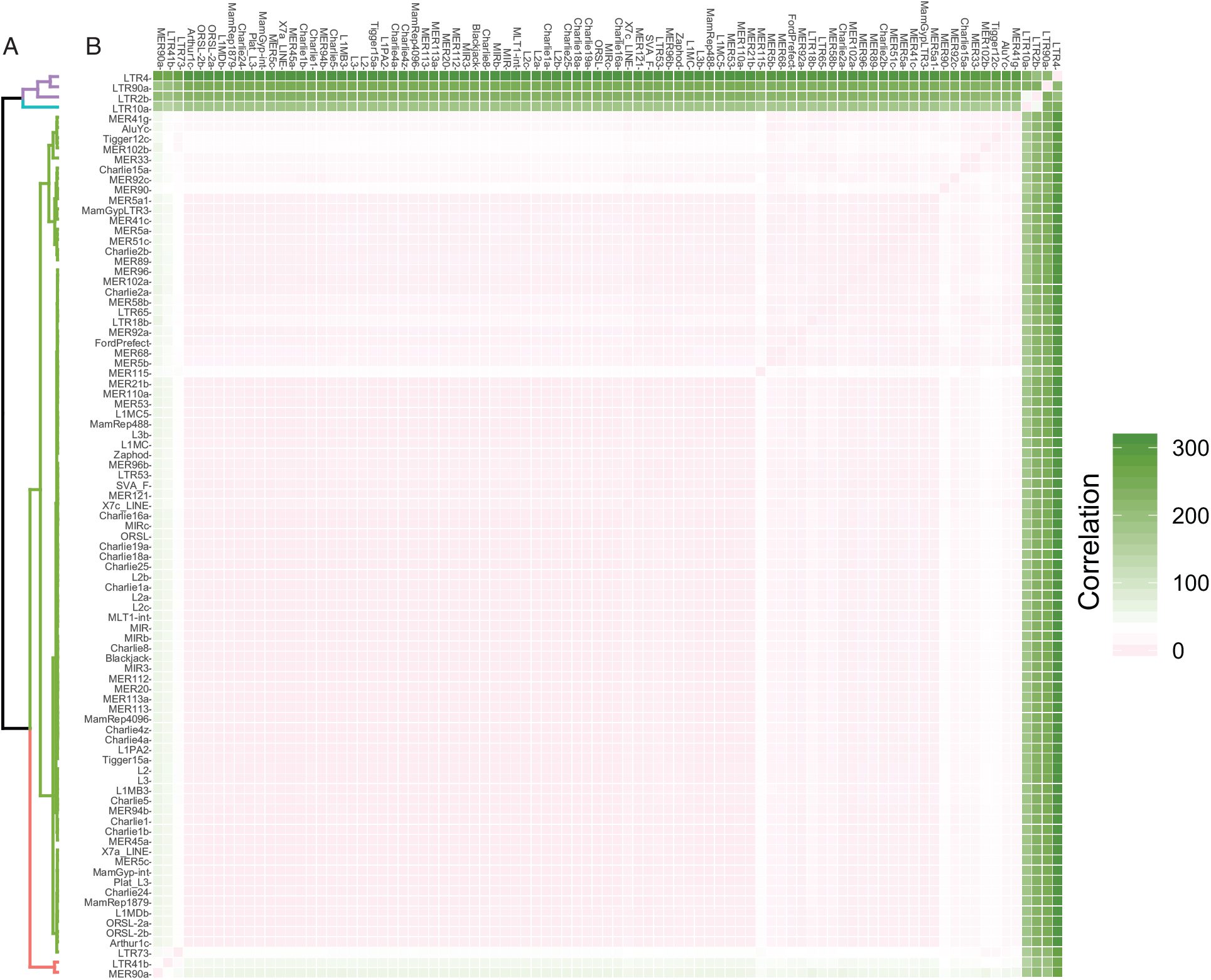
Hierarchical clustering of consensus TEs. (A) Hierarchical clustering (Manhattan distances) dendogram of consensus TEs based on mean luminescence values (log) and colored by K-means clustering (k=4). K-means clustering groups TEs into four categories corresponding to TEs with strong/moderate/weak enhancer or repressor functions. (B) Correlation matrix heatmap.

**Figure 5 – figure supplement 2.**
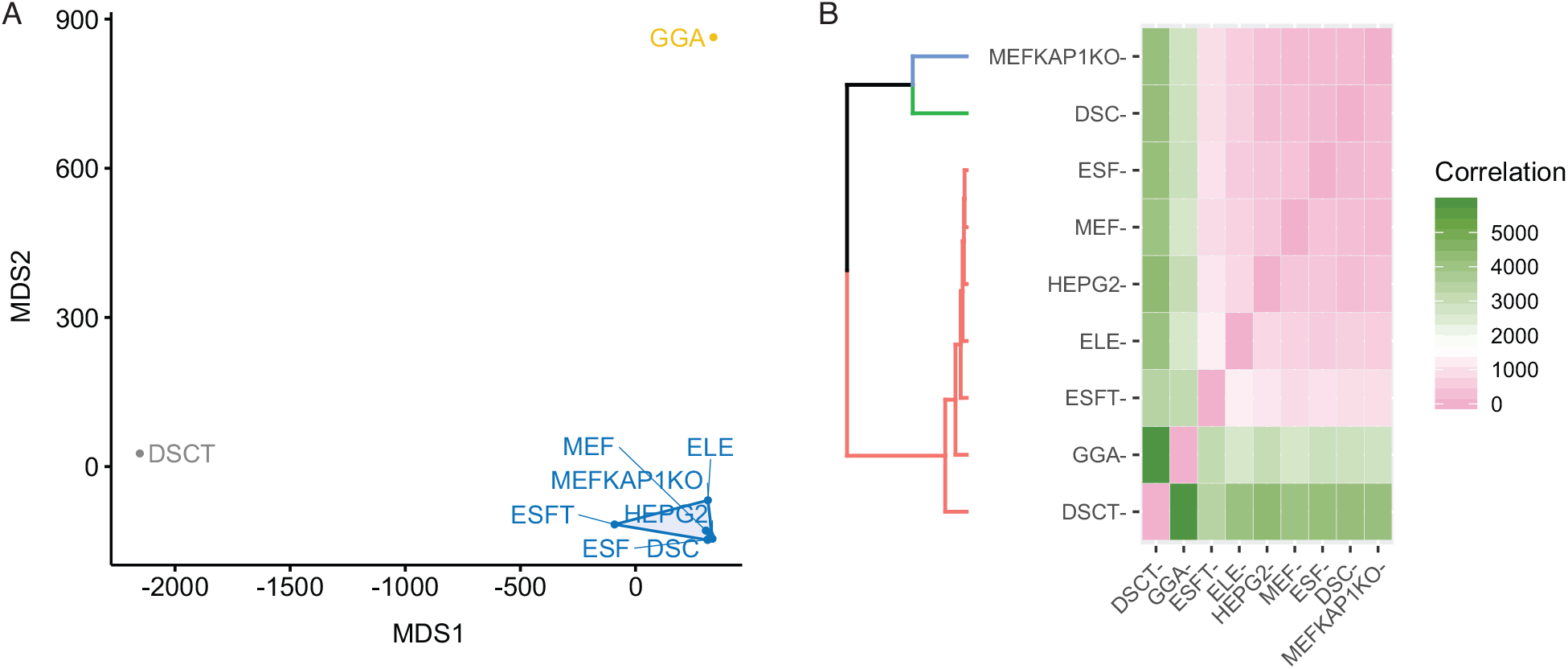
MDS and hierarchical clustering of cell types based on conTE luciferase results. (A) Multidimensional scaling plot (MDS) of cell-types based on consensus TE mean luminescence values (log) and grouped by K-means clustering (k=3). (B) Hierarchical clustering (Manhattan distances) dendogram and correlation matrix heatmap of cell-types based on consensus TE mean luminescence values (log) colored by K-means clustering (k=3).

**Figure 5 — Figure supplement 3.**
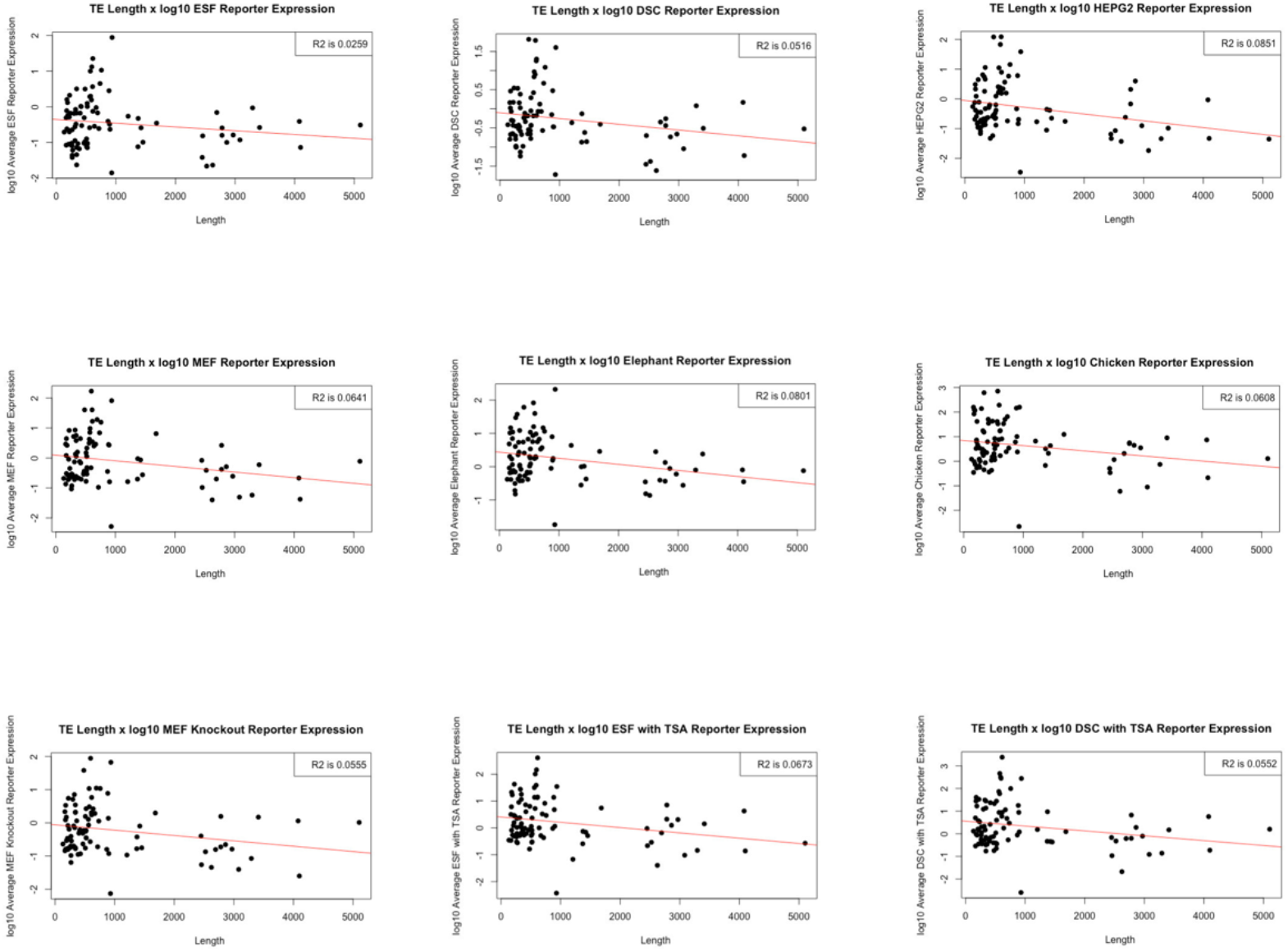
Consensus transposable elements (ConTEs) act as repressors independent of their length. Correlations between ConTE length and normalized luminescence (Log_10_) from dual luciferase reporter assays in each cell-type (ESF, DSC, HEPG2, MEF control and KAP1 KO, elephant fibroblasts, chicken fibroblasts) and treatment (ESF+TSA and DSC+TSA).

**Figure 5 — Figure supplement 4.** XSTREME motif discovery and enrichment analysis results with default settings.

**Figure 5 — Figure supplement 5.** XSTREME motif discovery and enrichment analysis results with default settings relaxed to allow for up to 15 motifs.

**Figure 5 — Figure supplement 6.** XSTREME motif discovery and enrichment analysis results specifically search for binding sites similar to PGR (NR3C2 binding site).

**Figure 5 — Source data 1. Consensus TE sequences.**

**Figure 5 — Source data 2. Dual luciferase assay results for six replicates in all cell-types.**

