## Supplementary figures and images for "Transposable elements continuously remodel the regulatory landscape, transcriptome, and function of decidual stromal cells"

### Cell HC.pdf

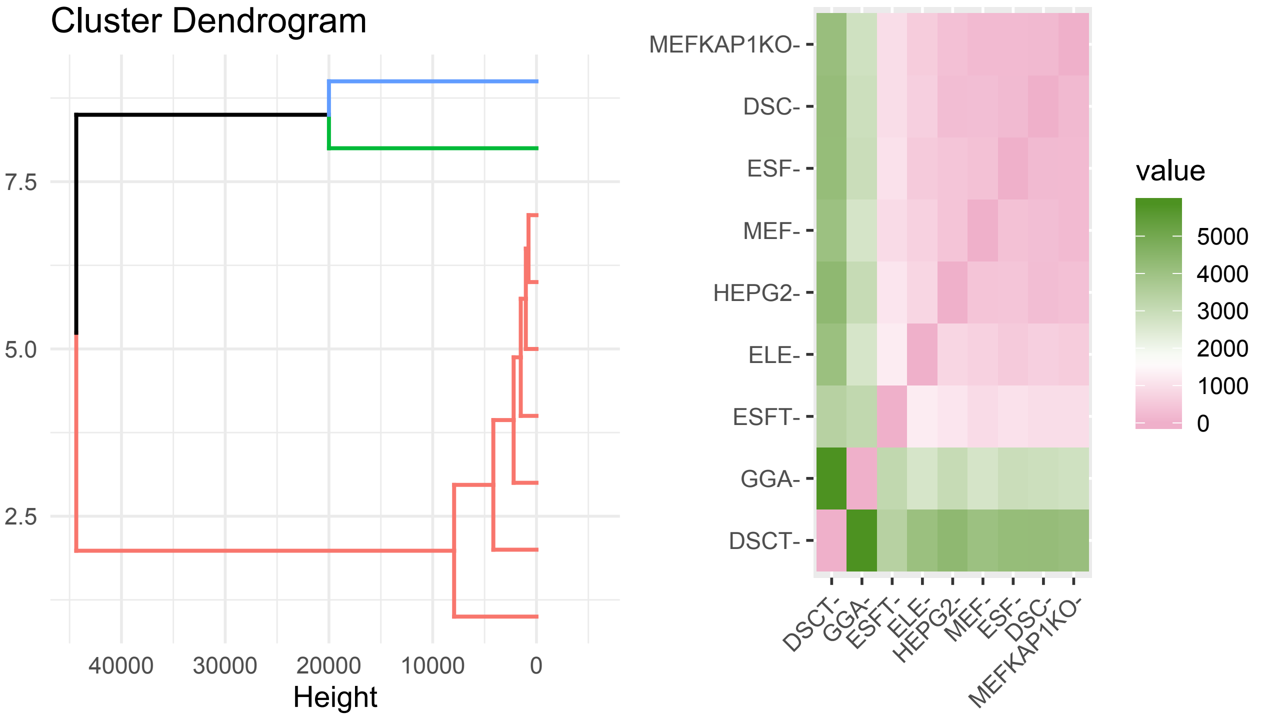

### HC.pdf

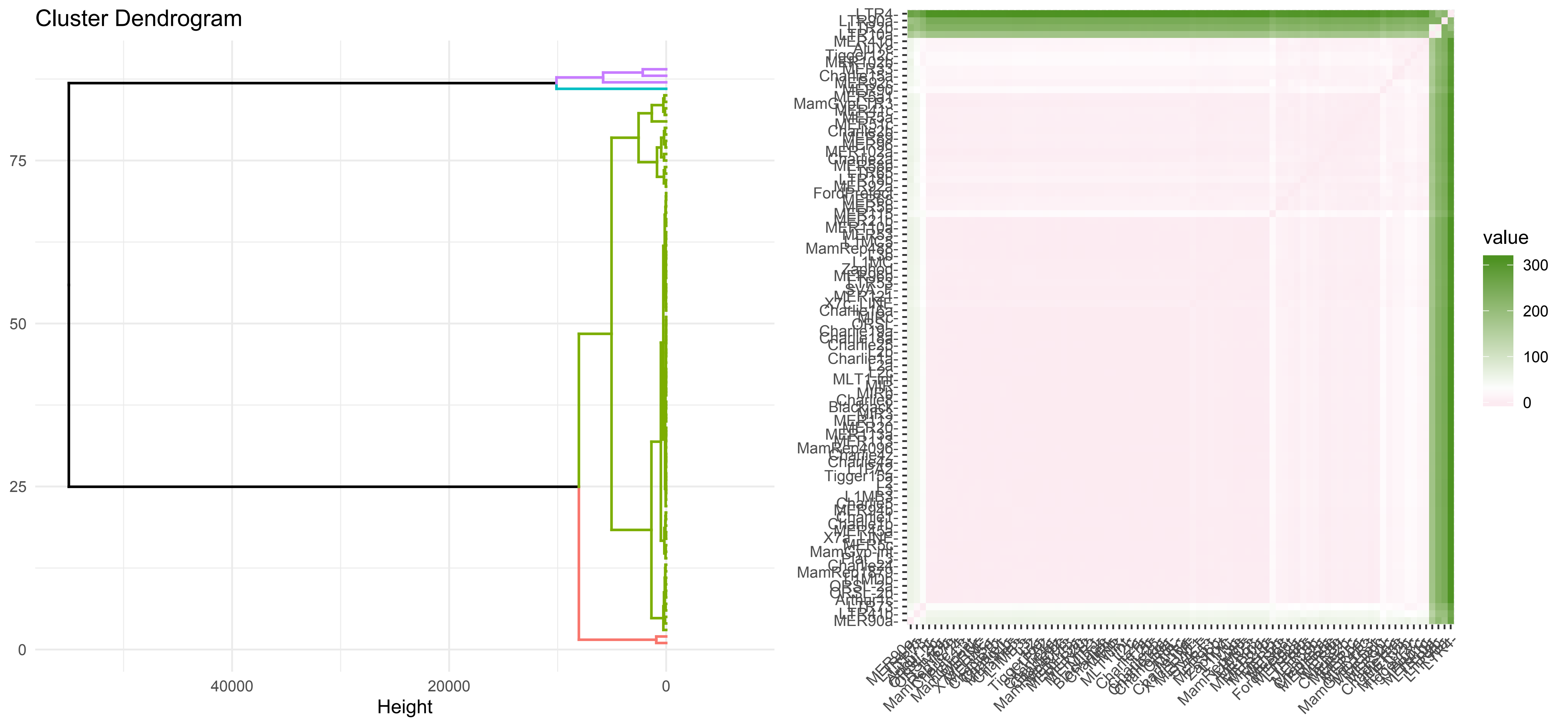

### MDS Cell.pdf

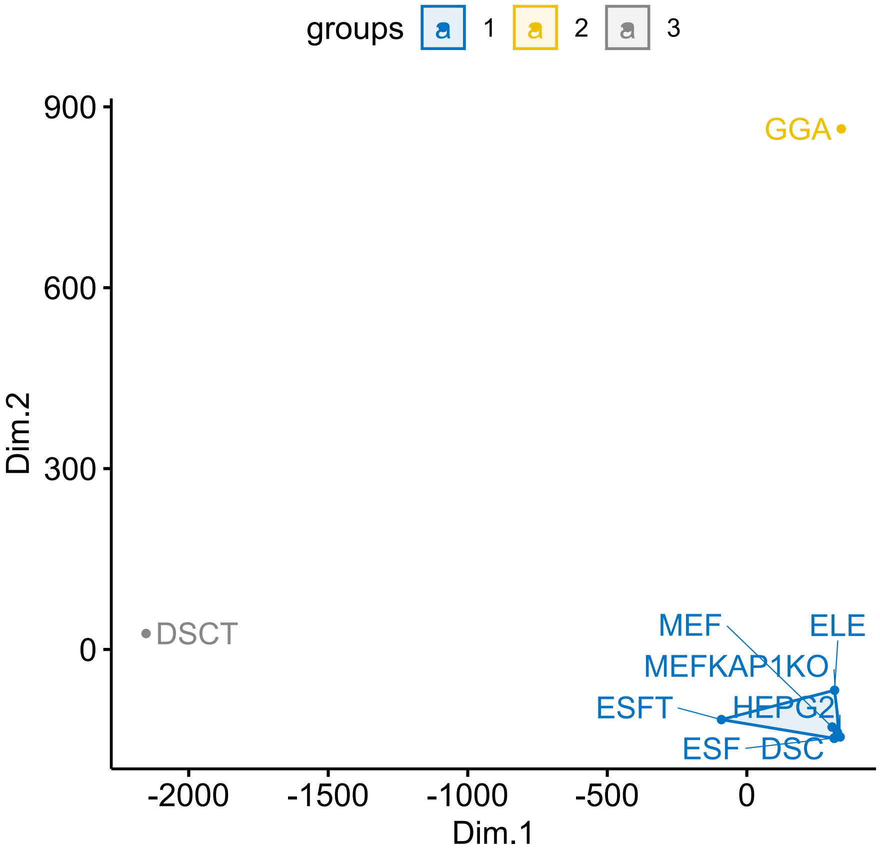

### MDS.pdf

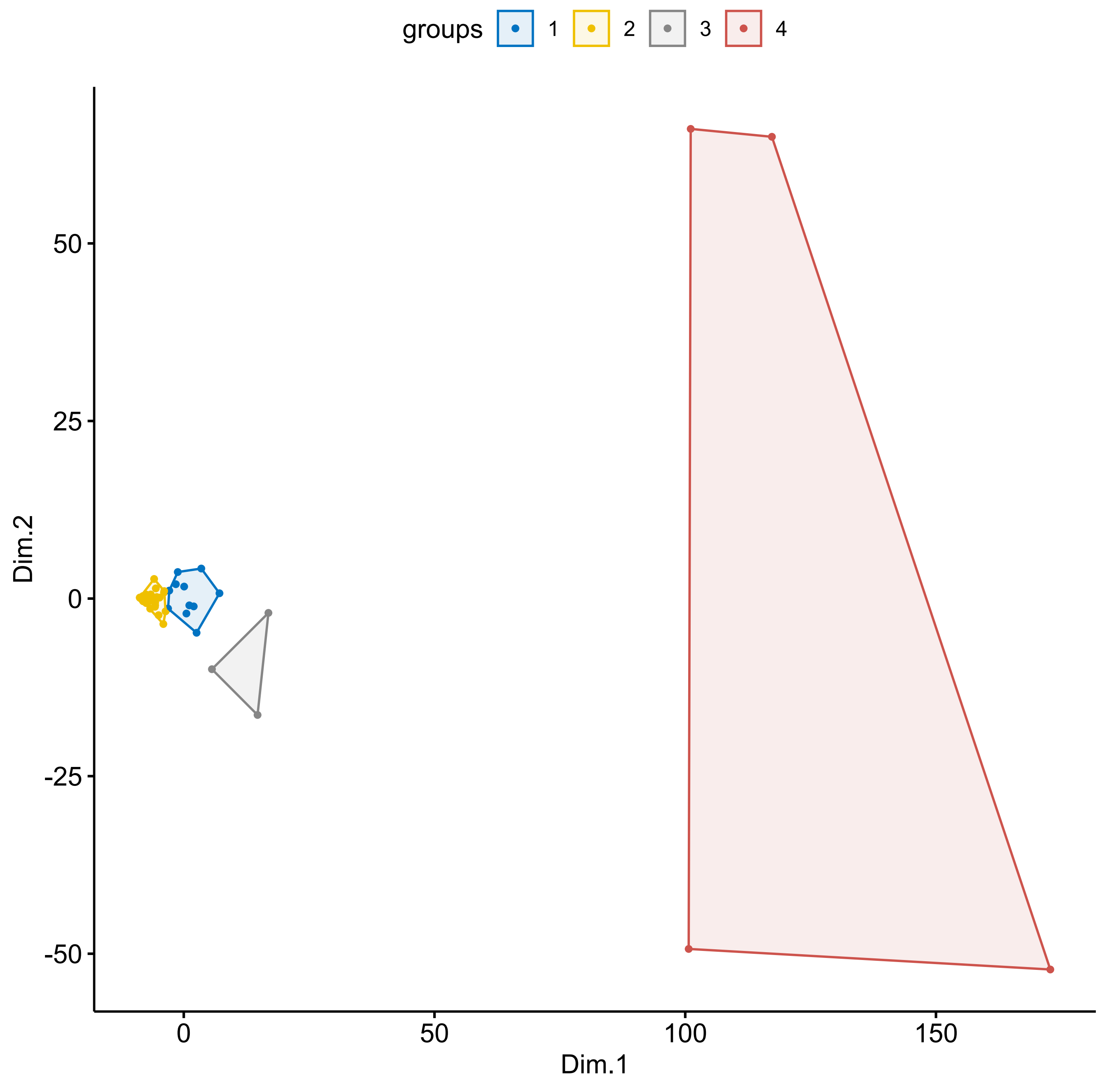
